# Hippocampal ripples signal contextually-mediated episodic recall

**DOI:** 10.1101/2021.06.07.447409

**Authors:** John J. Sakon, Michael J. Kahana

## Abstract

High-frequency oscillatory events, termed ripples, represent synchrony of neural activity in the brain. Recent evidence suggests medial temporal lobe (MTL) ripples support memory retrieval. However, it is unclear if ripples signal the reinstatement of episodic memories. Analyzing electrophysiological MTL recordings from 219 neurosurgical participants performing episodic recall tasks, we find that the rate of hippocampal ripples rises just prior to the free recall of recently-formed memories. This pre-recall ripple effect (PRE) is stronger in the CA1 and dentate gyrus (DG) subfields of hippocampus than neighboring MTL regions entorhinal and parahippocampal cortex. The PRE is also stronger prior to the retrieval of temporally and semantically clustered as compared with unclustered recalls, indicating the involvement of ripples in contextual reinstatement, which is a hallmark of episodic memory.

**One sentence summary:** High frequency human hippocampal ripples occur prior to those memories most likely retrieved via episodic mechanisms.

---

Experiments in animal models have linked high frequency ripples to consolidation and replay during quiescent and sleep states (*1*), and more recent work has linked ripples to awake behavior (*2–5*). Converging evidence from these animal studies (*5*), computational modeling (*6*), and new examinations in humans support the hypothesis that hippocampal (*7–9*) or surrounding medial temporal lobe (MTL) (*10, 11*) ripples signal upcoming memory retrieval. But it is unknown if such ripples are always present during retrieval of recently-experienced stimuli (*7, 10*) or if they are specific to the reinstatement of contextual information (*12–14*), a key feature in models of memory retrieval (*12, 15*) shown to be critical for episodic memory retrieval in humans (*16*). In short, we ask if MTL ripples during awake behavior are specific to episodic memories.

We answer this question using two intracranial EEG datasets of participants with electrodes in hippocampal subfields CA1 or DG, regions critical for episodic memory (*17–19*), as well as neighboring entorhinal or parahippocampal cortex. Participants were tested on at least one of two memory paradigms: delayed free recall of unrelated word lists (FR, 180 participants, 739 bipolar electrode pairs, **Fig. 1a**) and delayed free recall of categorized word lists (catFR, 104 participants with 65 of them also FR participants, 373 bipolar electrode pairs, **Fig. 4a**). Free recall, in which participants study a list of sequentially presented items and subsequently attempt to recall them in any order, allows researchers to isolate the processes underlying episodic memory retrieval (*20*). Transitions between consecutively recalled items enable the identification of neural processes underlying contextual reinstatement as it relates to both semantic and temporal associations among studied items (*13, 14, 21*). By relating ripples to how these associations organize recall, as seen in the analysis of recall transitions, we aim to elucidate the relationship between ripples and contextually-mediated retrieval processes.

**Fig 1.**
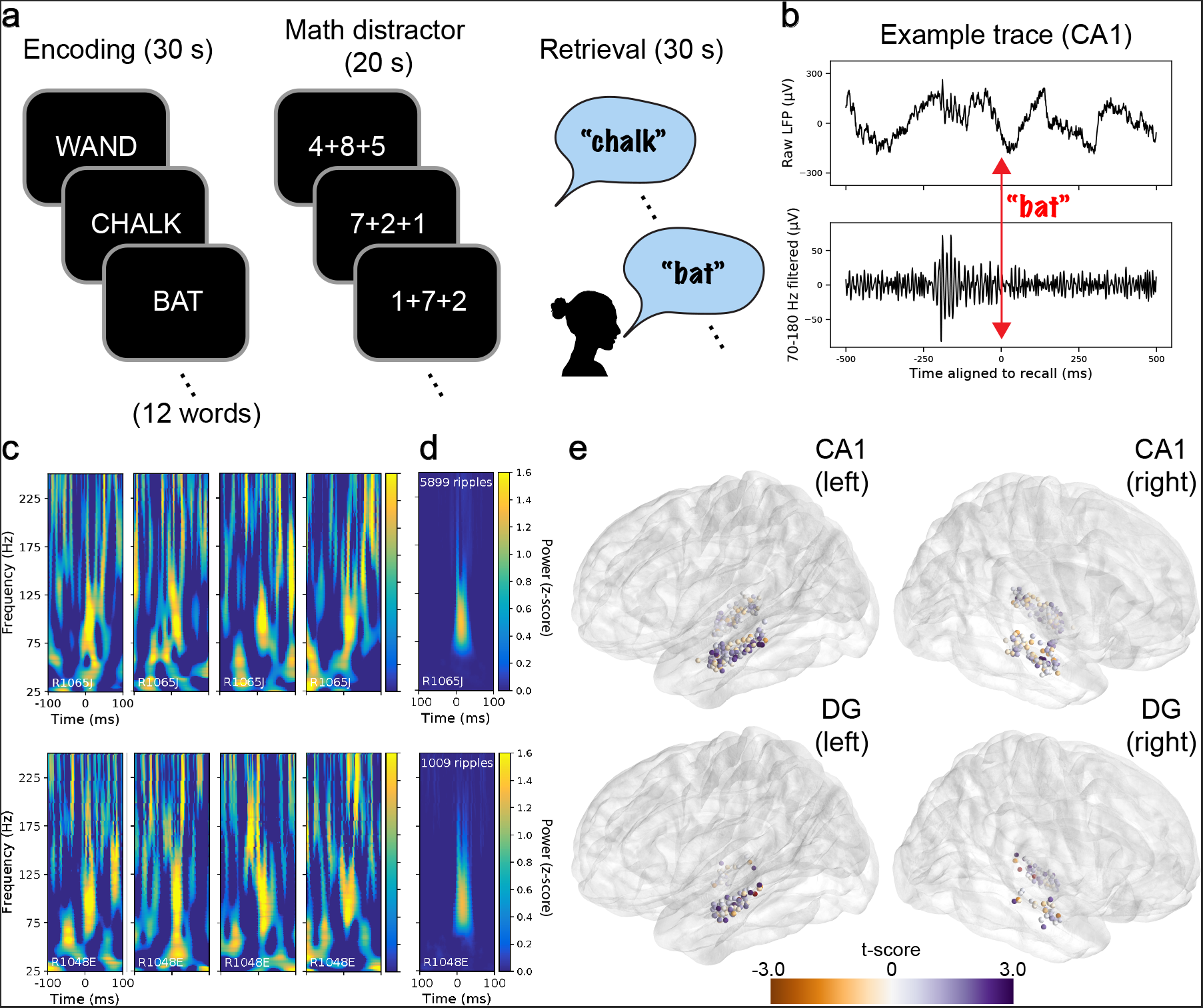
Free recall task and ripple details. **a**, Each list of the free recall (FR) task consists of an encoding period with 12 words presented sequentially, an arithmetic distractor, and a verbal free recall phase. **b**, Example hippocampal CA1 trace of raw (top) and Hamming bandpass-filtered (bottom) local field potential (LFP) aligned to the time of recall vocalization. Red indicates alignment to time of recall when the participant says “bat”. **c**, Example spectrograms of single ripples detected in CA1 for two participants, 4 from each. Each plot shows 100 ms before and after aligned to the start of a single ripple event. **d**, Average spectrograms for all ripples across sessions in CA1 for the same two participants. **e**, Localization of hippocampal CA1 and dentate gyrus (DG) electrode pairs for all FR participants. Views are all sagittal with ~10° axial tilt so both hippocampi are visible in each plot. Electrode pairs are colorcoded by t-scores of pre-recall ripple rise **(Eq. 2)**, with purpler colors indicating a stronger rise in ripples before recall. CA1, N=205; DG, N=100.

To detect ripples we use an algorithm recently shown to isolate such high-frequency events in human hippocampus during both memory encoding and retrieval (*7*) (Methods). Ripple peak frequencies (**Fig. 1c**), durations, spatial proximity, and rates (**Supplementary Figs. 1-2**) are similar to previous work (*7, 8, 10*). Anatomical localization of electrodes was performed by a combination of neuroradiologist labels and automated segmentation via separate processes for hippocampal subfields (*22*) and entorhinal and parahippocampal cortices (*23, 24*).

We partitioned our data into two halves: a first half for developing initial analyses, and a second half held out as a confirmatory dataset. We pre-registered our hypotheses as well as the initial figures for the first half of the data on the Open Science Framework (https://osf.io/y5zwt). Therefore, for the main tests throughout the manuscript, we present two sets of statistics: 1) the significance of model coefficients on the held out half of data (the “ held out” data) and 2) the significance of model coefficients on the full dataset (Methods).

The analyses detail three main findings. First, we establish the pre-recall ripple effect, in which ripples occur just prior to recalls that are not the first recalled from each list. Next, we find this effect is strongest in hippocampal subfields CA1 and DG. Finally, we show that the pre-recall ripple effect is strongest on trials that reinstate episodic information.

To elucidate the relation between ripples and recall we align hippocampal recordings to the onset of each correct recall vocalization in the FR dataset. A raster plot for 10 example participants with hippocampal recordings illustrates when ripples occur with respect to these recalls, where each row is a recall and each dot represents the start time of a single ripple (**Fig. 2a**). The raster suggests that ripple rates rise several hundred ms prior to vocalization onset, as shown in recent work (*7, 8, 10*).

**Fig 2.**
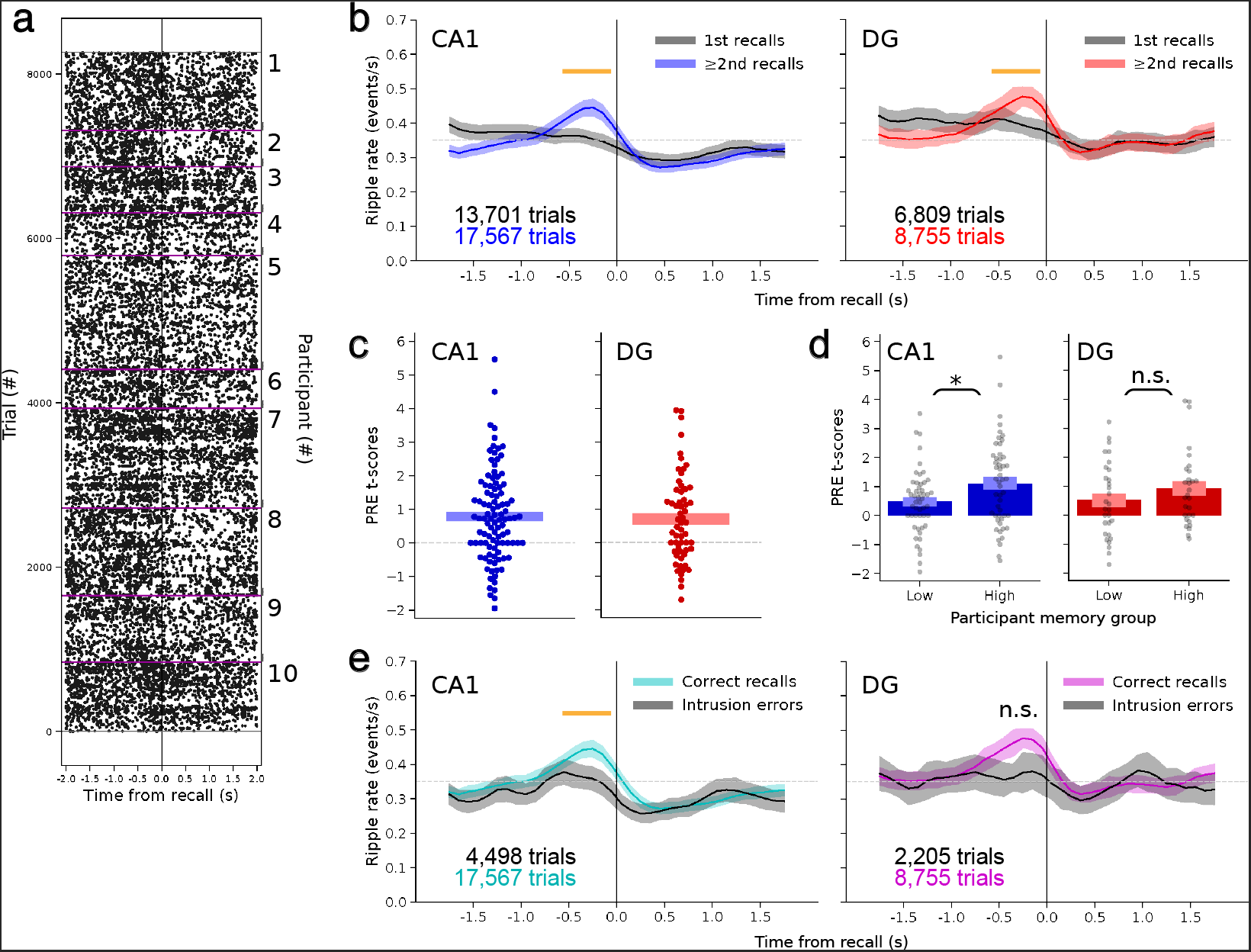
High frequency ripples increase in hippocampal subfields CA1 and dentate gyrus shortly before free recall. **a**, Raster plot aligned to free recall for all hippocampally-localized electrode pairs in 10 selected participants. Each dot represents the start time of a ripple. Purple lines divide participants. **b**, Peri-vocalization time histograms (PVTHs) for hippocampal subfields CA1 and dentate gyrus (DG) separated by whether the recall was the 1st made during the retrieval period or ≥2nd during the retrieval period for FR dataset. Trial numbers for each are labeled, where a trial is defined as a single recall on a single electrode pair. Significance of interaction from mixed model assessing the **p**re-recall **r**ipple **e**ffect (**PRE**) for ≥2nd recalls (Eq. 1): held out data: CA1, *β* = 0.12±0.023, *P* = 3.9 × 10^−6^, DG, *β* = 0.091±0.033, *P* = 0.010; 100% of data: CA1, *β* = 0.11±0.025, *P* = 2.4 × 10^−5^, DG, *β* = 0.10±0.025, *P* = 1.6 × 10^−4^ (FDR-corrected across 6 tests of Eq. 1 across **Figs. 2-4**). CA1: 263 sessions from 131 participants. DG: 171 sessions from 83 participants. Error bands are SE from a separate mixed model calculated at each time bin. Dotted gray line is to aid in visual comparison between PVTHs throughout the paper. Orange line indicates significant time range. **c**, t-scores for each participant from a mixed model assessing **PRE** for ≥2nd recalls for areas CA1 and DG (Eq. 2). Positive values indicate a stronger **PRE**. Only participants with ≥20 recalls are included (CA1: N=100; DG: 59). Bars indicate ±1 standard error from mean. One-sample t-test of t-scores from Eq. 2 vs. 0: held out data: CA1, *P* = 3.9 × 10^−4^, t = 4.1, df = 58; DG, *P* = 2.7 × 10^−3^, t = 3.5, df = 37; 100% of data: CA1, *P* = 3.3 × 10^−7^, t = 5.9, df = 99; DG, *P* = 1.9 × 10^−4^, t = 4.2, df = 58 (FDR-corrected for 6 t-tests across **Figs. 2-4**). **d**, t-scores for even split of participants based on their average recalls per list (each participant in gray). Bars are mean (dark) ± standard error (light). T-test of t-scores between the low and high recall participants: CA1, *P* = 0.036, t = 2.4, df = 98; DG, *P* = 0.23, t = 1.2, df = 57 (FDR-corrected across 2 tests). Significance of the interaction between rise in **PRE** and number of recalls per list is assessed via a mixed model (Eq. 3): CA1, *β* = 0.028±0.009, *P* = 0.0046; DG, *β* = 0.011±0.014, *P* = 0.45 (FDR-corrected across 2 tests). Same participant N as **c**. Error bands are SEs. **e**, PVTHs for correct ≥2nd recalls vs. ≥2nd intrusions. Conventions and participant N are the same as **b**. Significance of **PRE** for correct vs. intrusion trials (Eq. 5): held out data: CA1, *β* = −0.081±0.026, *P* = 0.0037, DG, *β* = −0.038±0.036, *P* = 0.30; 100% of data: CA1, *β* = −0.067±0.021, *P* = 0.0028, DG, *β* = −0.043±0.027, *P* = 0.11 (FDR-corrected across 2 tests).

Models of free recall posit separate mechanisms for recall initiation and subsequent retrieval transitions, with the former being driven by a persistent representation of items or context, and the latter being driven by cue-dependent associative retrieval (*20, 25*). Recordings from hippocampal subfields CA1 and DG, averaged across all participants into peri-vocalization time histograms (PVTHs), reveal clear physiological evidence for this distinction. Specifically, cue-dependent recalls (i.e., those following the first response, or ≥2nd recalls) exhibit a sharp pre-retrieval rise in ripples (**Fig. 2b**), which we term the **p**re-recall **r**ipple **e**ffect (**PRE** for the remainder of the paper). In contrast, the 1st recall in each retrieval period does not show this same **PRE** (**Fig. 2b**).

Using a linear mixed effects model to quantify this distinction while accounting for both within and between participant variability **(Eq. 1)**, the **PRE** is significantly stronger for ≥2nd recalls compared to 1st recalls in both CA1 (**Fig. 2b**, left) and DG (**Fig. 2b**, right). Further, the **PRE** is significant across participants when looking at only ≥2nd recalls for both CA1 and DG (**Fig. 2c; Eq. 2**). The localization of each bipolar electrode pair to the CA1 or DG hippocampal subfields is taken as the midpoint between adjacent electrode contacts. However, if either of the two contacts are outside the subfield, the ripples for this pair could possibly originate from a different region. To prove ripples indeed originate from both CA1 and DG, we performed a control analysis using only the CA1 pairs where both contacts themselves were localized to CA1, and only the DG pairs where both contacts themselves were localized to DG. Even with these more conservatively selected pairs, we find a significant PRE with similar ripple rates for both subfields (Supplementary Fig. 3).

As the first recall on each list tends to happen early in the retrieval period, is the absence of a **PRE** for these trials related to the absolute time of recall, and not the order of 1st v. 2nd? Looking at only those recalls that occur within the first 5.0 s of the retrieval period, the **PRE** remains significantly stronger for ≥2nd recalls than 1st recalls (**Supplementary Fig. 4a**), suggesting the **PRE** is correlated with recalls that follow a previous recall (*26*) and not just recalls occurring later in the retrieval period. Meanwhile, when the 1st recall does occur later in the retrieval period (after 5.0 s), the **PRE** is no different between 1st and ≥2nd recalls (**Supplementary Fig. 4b**), further evidence that ripples are linked to the processes governing retrieval and not persistent representations held from encoding (*27*) or retrieved just prior to the start of the free recall period.

Does a participant’s strength of the **PRE**, as measured for ≥2nd recalls, correlate with memory performance? Splitting the participants from the FR dataset into those with the highest and lowest number of recalls per list and comparing the **PRE** between these groups, participants with high recall show a significantly stronger **PRE** in CA1 and in the same direction for DG (**Fig. 2d**). Using a linear mixed model to quantify the relation between each participant’s list-level recall performance and their rise in ripple rate from the **PRE (Eq. 3)** further evidences this effect, as there is a significant positive interaction between participant memory and the **PRE** for CA1 electrode pairs and in the same direction for DG pairs (**Fig. 2d caption**). In addition, comparing correct recalls with intrusions (i.e. recalls of items not present on the target list (*28*)) reveals a significantly stronger **PRE** for correct recalls in CA1 and in the same direction for DG (**Fig. 2e**). Taken together, the link between the **PRE** and correct, cue-dependent recall implicates hippocampal ripples in episodic memory retrieval.

The frequency range for the ripple detection algorithm—based on a recent study of human hippocampal ripples (*7*)—is relatively broad (70-178 Hz). This range likely includes sharp-wave ripple–associated fast-gamma as well as ripples (*29, 30*). Whereas previous work has grouped these events as they differ only in frequency and relative amplitudes between subfields (*29, 30*), we ask if ripples detected using algorithms with narrower ranges still reliably show a **PRE**. For a first check, we implement a ripple detection algorithm with a narrower range (80-120 Hz) that was recently used to identify ripples in MTL (*8,10,11*). This stricter algorithm yields lower ripple rates with a wider distribution of durations and a peak ripple frequency ~90 Hz (**Supplementary Figs. 1c&5a**), all similar to previous work (*8, 10, 11*). Despite the lower ripple rates, the **PRE** is significant for ≥2nd recalls compared to 1st recalls in CA1 and in the same direction for DG (**Supplementary Fig. 5b**). For a second check, we utilize the original ripple detection algorithm, but with a higher frequency range (125-200 Hz) to isolate ripples at frequencies typically reported in rodent sharp-wave ripple work (*1,29*). This method once again yields lower ripple rates but with a similar distribution of durations as the original algorithm and a frequency peak ~150 Hz (**Supplementary Figs. 1b&6a**). Once again, we confirm the main result as the **PRE** is significant for ≥2nd recalls in both CA1 and DG (**Supplementary Fig. 6b**).

Finally, we address the possibility that **PRE** is related to seizurogenic tissue in epileptic participants, even though recent work suggests epileptiform tissue should show a weaker link between ripples and memory than healthy tissue (*8*). For those participants with a clinically-defined seizure onset zone (SOZ), we take all trials from bipolar pairs in the SOZ and compare them to all trials from bipolar pairs not in the SOZ. For each hippocampal subfield CA1 and DG, both SOZ and non-SOZ trials show a significant **PRE**; however, neither subfield shows a significant difference when comparing the **PRE** between them (**Supplementary Fig. 7a**), suggesting **PRE** is not related to epileptic activity.

In addition to hippocampal electrode pairs, many participants had electrode coverage in entorhinal and parahippocampal cortex (**Fig. 3a-b**). Ripples are known to occur in both these regions (*1, 10*), so we asked if a **PRE** occurs before recalls in the FR dataset as shown in hippocampus. Indeed, entorhinal cortex shows a significant interaction between ≥2nd recalls and **PRE** when averaging recordings across all participants (**Fig. 3c, Eq. 1**), as well as significant t-scores at the participant-level for ≥2nd recalls (**Fig. 3e**). Parahippocampal cortex ripples were not significant for either of these tests (**Fig. 3d & 3f**).

**Fig 3.**
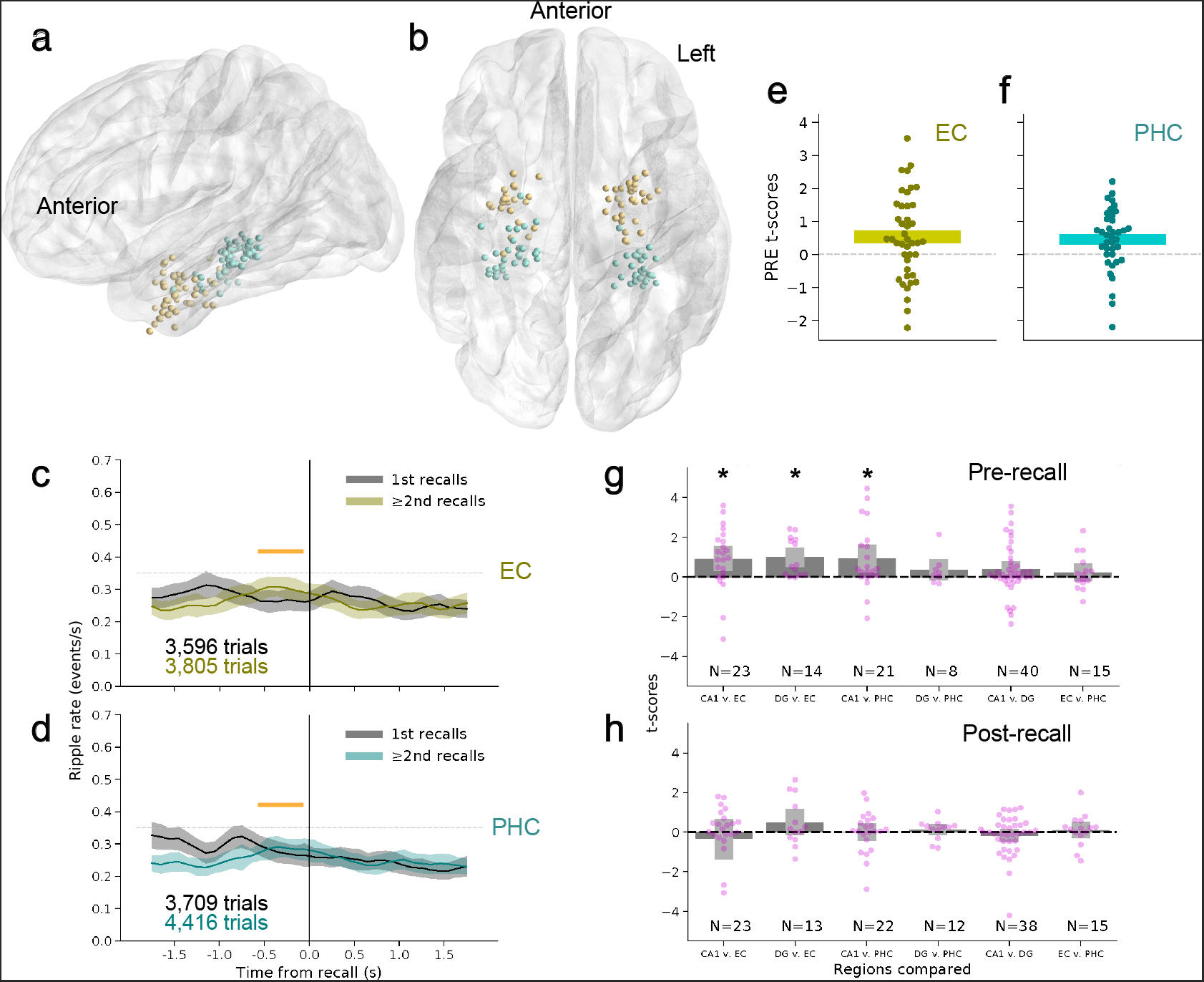
Regional differences in the pre-recall ripple effect (PRE). **a**, Localization of extrahippocampal electrode pairs from left sagittal perspective. Entorhinal, N=53; Parahippocampal, N=61. **b**, Ventral perspective. **c**, PVTH for entorhinal cortex trials, separated by 1st recall and ≥2nd recalls from each list for FR dataset. Significance of mixed model assessing **PRE** for ≥2nd recalls (Eq. 1): held out data: *β* = 0.091±0.035, *P* = 0.014, 100% of data: *β* = 0.093±0.03, *P* = 0.0029 (FDR-corrected across 6 tests of Eq. 1). 127 sessions from 70 participants. Error bands are SE from a mixed model at each bin and orange line indicates significant time range as in Fig. 2. **d**, Same for parahippocampal cortex. 118 sessions from 63 participants. Held out data: *β* = 0.046±0.031, *P* = 0.15, 100% of data: *β* = 0.071±0.028, *P* = 0.013. **e**, Same conventions as **Fig. 2c**, with mixed model t-scores for **PRE** calculated for entorhinal electrode pairs in each participant (Eq. 2). N=42 participants. Held out data: *P* = 0.054, t = 2.2, df = 31; 100% of data: *P* = 0.012, t = 2.7, df = 41 (FDR-corrected across 6 tests). **f**, Same as **e** but for parahippocampal cortex. N=38 participants. Held out data: *P* = 0.078, t = 1.9, df = 25; 100% of data: *P* = 0.0078, t = 3.0, df = 37 (FDR-corrected across 6 tests). **g**, Mixed model t-scores of pairwise comparisons of **PRE** for each participant with electrodes in at least two of the four regions under study: hippocampal areas CA1 and DG as well as entorhinal cortex (EC) and parahippocampal cortex (PHC). The model assesses **PRE** for ≥2nd recalls from a time range −600 to −100 ms before recall (Eq. 4). Asterisks indicate the first of the pair being compared is significantly greater than the second (*P<*0.05, FDR-corrected for 6 pairwise comparisons of pre-recall ripples using Eq. 4). Sample size of participants indicated at bottom of each comparison. **h**, Similar to **g**, but for a mixed model assessing a drop in ≥2nd recalls from a time range 200 to 700 ms after recall (Eq. 4). No comparison shows a significant drop in ripples after recall (*P<*0.05, FDR-corrected for 6 pairwise comparisons of post-recall ripples using Eq. 4). Error bars are SE throughout.

To directly compare the **PRE** between regions, we contrast them in a single model. We make pairwise comparisons between the hippocampal subfields (CA1 and DG) and entorhinal and parahippocampal cortices, but only for those participants with bipolar electrode pairs in at least two of these regions (e.g., a participant with electrodes in CA1, DG, and entorhinal cortex would contribute 3 pairwise comparisons). A separate linear mixed model for each participant, which accounts for differences between sessions with random effects, compares the **PRE** on ≥2nd trials between pairs of regions **(Eq. 4)**. The t-scores from this model are then combined for a one-sample t-test across participants (**Fig. 3g**). Both hippocampal subfields CA1 and DG have a significantly stronger **PRE** than entorhinal cortex. CA1 has a significantly stronger **PRE** than parahippocampal cortex, while DG vs. parahippocampal cortex is not significant, likely owing to having the smallest sample (N=8 participants). There are no reliable differences in the **PRE** between CA1 and DG or between entorhinal and parahippocampal cortex. We next asked whether the post-recall drop in ripple rate evident in many participants (Fig. 2a-b), possibly due to a refractory period after the rise in ripples from the **PRE** (*1, 31*), is also specific to hippocampus. Taking advantage of these same participants with electrode pairs in at least two regions, there is no evidence that any particular region drops in ripple rate after recall more than any other (**Fig. 3h**), suggesting the post-recall drop is not consistent like the **PRE**.

Finally, we ask if the **PRE** correlates with behavioral measures specific to episodic memory (*28, 32, 33*). We first focus on the catFR dataset as the list of words in this task has a rich semantic and temporal structure (**Fig. 4a**). In particular, words in the catFR task are drawn from a pool of 25 semantically-related categories with three categories selected per 12-word list. Each set of four words from a category are presented as pairs with the pairs never shown back-to-back. For example, dolphin and octopus might be a pair of consecutively shown words followed by cupcake and pie, which are then followed by fish and whale (**Fig. 4a**). This setup allows us to measure contextual reinstatement in semantic and temporal dimensions when participants recall the words, as back-to-back recalls can transition between 1) a semantic pair that was temporally adjacent in the list (20% of recalls), 2) a semantic pair that was temporally remote in the list (20% of recalls), and 3) a pair of words that were temporally adjacent in the list but not semantically-related (only 3% of recalls, as participants tend to recall via semantic associations in catFR, so we do not investigate them further). The remaining transitions are remote unclustered (17%), meaning two semantically unrelated words that were not adjacent on the list. By comparing groups of trials with contextual associations to those without, we can assess if ripples not only precede recall, but also precede reinstatement of contextual information used to remember items, a key signature of episodic memory (*12*). Note that we only use ≥2nd recalls in these analyses, as the 1st recall in every list does not show the signature **PRE**(**Fig. 2b & Fig. 4c-d**), likely due to weaker contextual reinstatement before the first recall (see **Discussion**).

**Fig 4.**
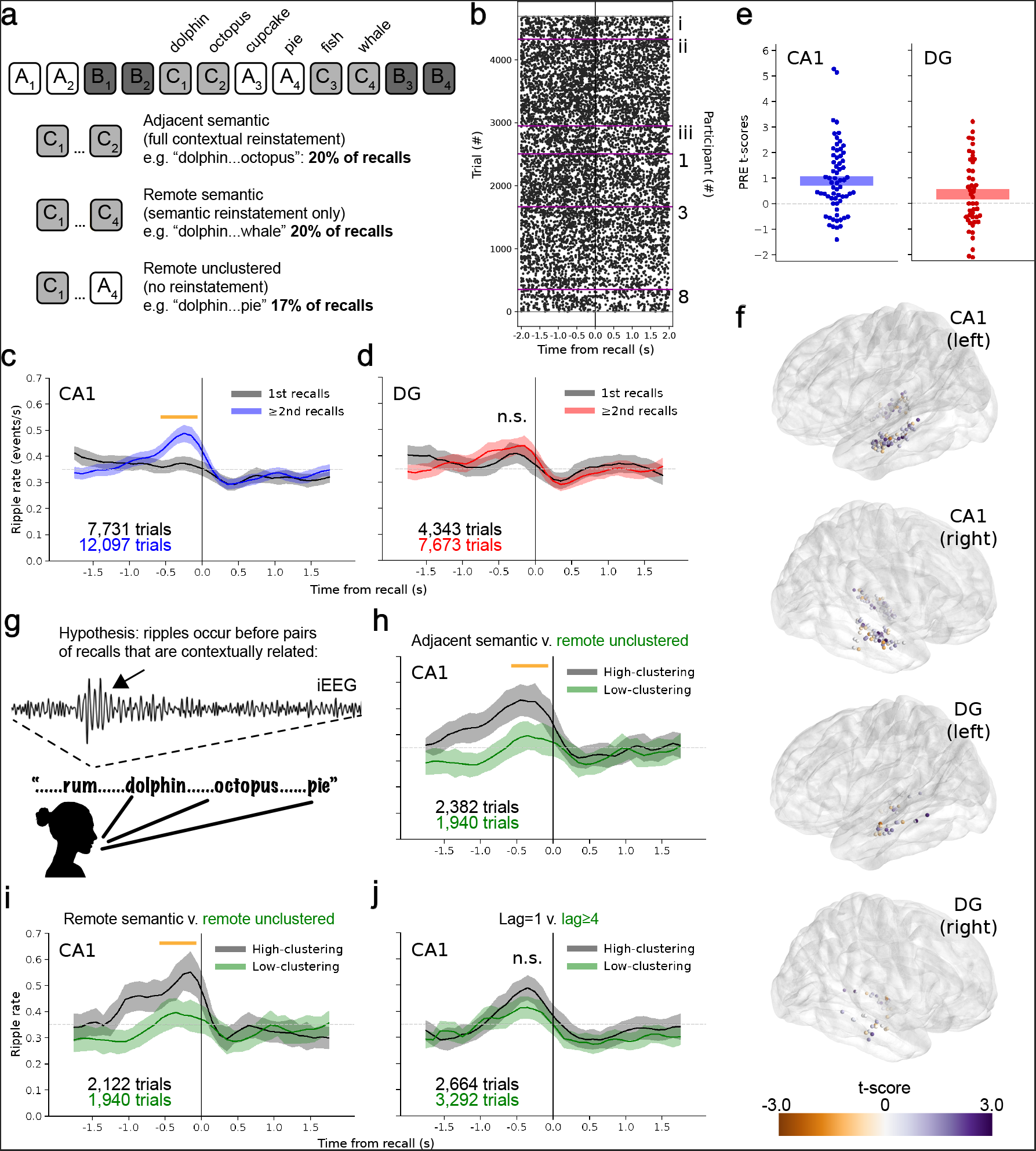
Context reinstatement and the pre-recall ripple effect (PRE). **a**, (top) Outline of categorized free recall task (catFR). Word lists were comprised of 12 words from 3 semantic categories (shown as A_x_, B_x_, and C_x_) and shown during encoding in pairs of two. (bottom) Percentages of recall types by transitions between recalls. **b**, Raster of ripples aligned to recall from three of the same participants in **Fig. 2a** that ran both task versions (1,3, and 8) and three new participants (i-iii). Purple lines divide participants. **c**, PVTH for hippocampal subfield CA1 aligned to recalls in catFR with same conventions as **Fig. 2b**. Significance of mixed model assessing **PRE** for ≥2nd recalls (Eq. 1): held out data: *β* = 0.12±0.029, *P* = 1.8 × 10^−4^, 100% of data: *β* = 0.12±0.022, *P* = 1.8 × 10^−7^ (FDR-corrected across 6 tests of Eq. 1). 177 sessions from 98 participants. Error bands are SE from a mixed model at each bin and orange line indicates significant time range as in Figs 2-3. **d**, Same for hippocampal area DG. Held out data: *β* = 0.068±0.048, *P* = 0.15, 100% of data: *β* = 0.046±0.032, *P* = 0.15. 124 sessions from 65 participants. **e**, Same conventions as **Fig. 2c**, with mixed model t-scores for **PRE** calculated for CA1 and DG electrode pairs in each participant performing catFR (Eq. 2). Held out data: CA1, *P* = 1.6 × 10^−4^, t = 4.7, df = 43; DG, *P* = 0.22, t = 1.25, df = 28; 100% of data: CA1, *P* = 1.7 × 10^−5^, t = 4.9, df = 65; DG, *P* = 0.077, t = 1.8, df = 45 (FDR-corrected across 6 tests). Only participants with ≥20 recalls are included (CA1: N=69; DG: 48). **f**, Localization of hippocampal electrode pairs in participants that ran catFR. Views are all sagittal with ~10° axial tilt so both hippocampi are visible in each plot. Electrode pairs are color-coded by t-scores of pre-recall rise, as in **Fig. 1e** (Eq. 2). CA1, N=136; DG, N=36. **g**, Schematic for hypothesis of ripples as a signature of contextual reinstatement. An example ripple before recall is shown (arrow) in zoomed-in iEEG (70-178 Hz filtered). **h**, PVTH of catFR trials comparing adjacent semantic vs. remote unclustered trials, a test of contextual reinstatement vs. no contextual reinstatement. Significance of coefficient comparing **PRE** for each trial type in mixed model (Eq. 5): held out data: CA1, *β* = −0.094±0.057, *P* = 0.19; 100% of data: CA1, *β* = −0.12±0.043, *P* = 0.027 (FDR-corrected across 6 tests of Eq. 5,, **Figs. 4h-j & Supplementary Fig. 8a-c**). Data from 145 sessions collected in 83 participants. Error bands are SE and orange line indicates significant time range as in Figs. 2-3. **i**, PVTH of catFR trials comparing remote semantic vs. remote unclustered trials, a test of semantic reinstatement vs. no contextual reinstatement. Same conventions, number of sessions and participants, and significance test as **h**. Held out data: CA1, *β* = −0.051±0.051, *P* = 0.47; 100% of data: CA1, *β* = −0.097±0.037, *P* = 0.027 (FDR-corrected). **j**, PVTH of FR data comparing adjacent recalls (lag = 1) vs. remote recalls (lag≥4), a test of temporal reinstatement. Data from 199 sessions collected in 109 participants. Same conventions and significance test as **h**. Held out data: CA1, *β* = −0.099±0.04, *P* = 0.084; 100% of data: CA1, *β* = −0.068±0.033, *P* = 0.078 (FDR-corrected).

Before assessing differences between types of recall, we confirm our main findings with the catFR dataset, which acts as an independent dataset to support our findings from the FR dataset. First, using three of the same participants that contributed to the FR raster plot and three new participants (**Fig. 2a**), a raster aligned to recall for the catFR task once again shows visual evidence of a **PRE** (**Fig. 4b**). Across all participants, the **PRE** is significant for ≥2nd recalls compared to 1st recalls in both CA1 (**Fig. 4c**) and DG (**Fig. 4d**). Looking at participants individually, there is a significant **PRE** for ≥2nd recalls across participants in CA1, and in the same direction for DG (**Fig. 4e**). However, due to randomness in participant electrode montages, there happened to be many fewer electrode pairs in DG than CA1 for catFR (36 vs. 136, respectively; **Fig. 4f**), in addition to fewer participants collected in catFR than FR (104 vs. 180, respectively), making this test relatively underpowered.

For the first test of contextual reinstatement, we set up a comparison between those recalls that act as the strongest contextual cues compared to those that act as the weakest. In particular, adjacent semantic (**Fig. 4a**), where the subsequent recall was both temporally adjacent and semantically related to the previous recall on the list, vs. remote unclustered, where the subsequent recall was neither. The hypothesis is that if ripples are a signature of contextual reinstatement, we expect a **PRE** *before* recall of the adjacent semantic pair (**Fig. 4g**). In other words, if one recall leads to a subsequent recall that is contextually associated with it, the expectation is the **PRE** before the initial recall is a signature of the reinstatement that leads to the transition from one recall to the next (*14, 34*). Indeed, in CA1, the **PRE** for adjacent semantic trials is significantly stronger than for remote unclustered trials (**Fig. 4h**). DG also shows a significantly greater **PRE** for adjacent semantic trials for this comparison (**Supplementary Fig. 8a**).

The next test of contextual reinstatement is a comparison between remote semantic and remote unclustered (**Fig. 4a**). This comparison isolates semantically-driven transitions, as all pairs of recalled words that were adjacent on the list are excluded. Once again, in CA1, there is a significantly greater **PRE** for remote semantic trials (**Fig. 4i**), although there is not a significant difference for these groups in DG (**Supplementary Fig. 8b**).

For the final test of contextual reinstatement, we aim to isolate temporal clustering based on the presentation order of the word list. Since the catFR task is designed to promote semantic associations, we return to the FR task, where the 12 words are not designed to be semantically related (**Fig 1a**). To assess temporal clustering we grouped all recalls that led to adjacent transitions from the list (absolute lag=1, 16% of transitions) and compare them to all recalls that led to remote transitions on the list (absolute lag ≥4, 20%) (*35*). The hypothesis remains the same: that the **PRE** should occur before those recalls leading to contextual reinstatement, in this case via temporal associations. For CA1, recalls that led to temporally clustered transitions show a stronger **PRE** than recalls that led to remote transitions, although this effect is not significant across multiple comparisons in Figs. 4h-j (**Fig. 4j**). There is no difference for DG trials (**Supplementary Fig. 8c**).

We investigated high frequency ripples as participants studied and subsequently recalled lists of unrelated items (N=180) or lists of categorically-related items (N=104). In both paradigms we found a punctate rise in ripples immediately before participants recall words. This pre-recall ripple effect (**PRE**) is specific to recalls that follow previously recalled items, signaling a cue-dependent retrieval process (**Fig. 2b & 4c-d**). Further, we find a stronger **PRE** for contextually-reinstated recalls (**Fig. 4h-j**). The **PRE** is also strongest in hippocampal subfields CA1 and DG compared to entorhinal and parahippocampal cortex (**Fig. 3g**). These results implicate ripples in hippocampally-initiated episodic memory retrieval.

The free recall task provides a window into the organization of memory because it permits people to report studied items in the order that they come to mind. The order and timing of recalled items reveals the temporal and semantic organization of memory, as people tend to consecutively recall temporally-proximate or semantically-related items (*36*). Modeling these dynamics of memory search has highlighted the importance of context–a latent representation that includes information about time, space, and semantics of recently experienced or recalled items (*20*). A key feature of these models is that remembering an item retrieves its prior contexts, which in turn triggers the next item that comes to mind. However, the first response is governed by the persistent context from the end of the list rather than the retrieved context caused by the preceding item (*25, 27*). Here, we find a stark dichotomy between the first recall on each list and subsequent recalls, with the **PRE** specifically occurring before subsequent recalls (**Figs. 2b & 4c-d**), suggesting that hippocampal ripples are a physiological correlate for retrieved context. The clustering results further ballast the link between hippocampal ripples and contextual reinstatement, as recalls with strong semantic and/or temporal association to the next recalled word show a significantly stronger **PRE** compared to recalls with low clustering (**Fig. 4g-j**).

In sum, hippocampal ripples preferentially occur before those recalls most likely to be achieved via contextual reinstatement of episodic memories, supporting the hypothesis that ripples mediate episodic memory retrieval (*1, 5*). Modeling results indicate amnesia in patients with MTL damage comes from an inability to reinstate context (*16*), which considered with the results presented here suggests a link between memory loss and ripple malfunction.

## Data and code availability

Data were collected as part of the DARPA Restoring Active Memory (RAM) initiative and is available to the public: http://memory.psych.upenn.edu/ElectrophysiologicalData. Code and processed data for all plots and analyses are available at: http://memory.psych.upenn.edu/files/pubs/SakoKaha21.code.tgz. Questions should be addressed to.

## Acknowledgements

Data was collected as part of the DARPA Restoring Active Memory (RAM) program (Cooperative Agreement N66001-14-2-4032). This work is supported by National Institutes of Health grant R01NS106611-02 and USAMRDC MTEC grant MTEC-20-06-MOM-013. The views, opinions, and/or findings contained in this material are those of the authors and should not be interpreted as representing the official views or policies of the Department of Defense or the U.S. Government. We also thank the Michael Kahana lab, the Joshua Jacobs lab, the

György Buzsáki lab, the Anli Liu lab, the Ehren Newman lab, Alice Healy, Dan Levenstein, and Sam McKenzie for providing valuable feedback on this work.

## Author contributions

J.J.S. analyzed data. J.J.S. and M.J.K. designed the study and wrote the paper.

## Supplementary Materials

Materials and Methods

Fig S1-S8

## Materials and Methods

### Human participants

Our dataset included 219 adult participants in the hospital for medication-resistant epilepsy with subdural electrodes placed on the cortical surface or within the brain for the purpose of localizing epileptic activity. Data were recorded at 8 hospitals from 2015-2021. These include Thomas Jefferson University Hospital (Philadelphia, PA), University of Texas Southwestern Medical Center (Dallas, TX), Emory University Hospital (Atlanta, GA), Dartmouth-Hitchcock Medical Center (Lebanon, NH), Hospital of the University of Pennsylvania (Philadelphia, PA), Mayo Clinic (Rochester, MN), National Institutes of Health (Bethesda, MD), and Columbia University Hospital (New York, NY). Before data was collected at any hospital our research protocol was approved by the Institutional Review Board at the University of Pennsylvania via a reliance agreement or at participating hospitals.

### Free recall task (FR)

participants performed a delayed free recall task where for each “list” they viewed a sequence of common nouns with the intention to commit them to memory. The task was run at bedside on a laptop and participants were tasked to finish up to 25 lists for a whole session or 12 lists for a half-session. The free recall task consisted of four phases per list: countdown, encoding, distractor, and retrieval (**Fig. 1a**). Each list began with a 10-second countdown period with numbers displayed on the screen from 10 to 1. Next was encoding, where participants were sequentially presented a list of 12 words centered on the screen that were selected at random–without replacement in each whole session or two consecutive half sessions–from a pool of 300 high frequency, intermediate-memorable English or Spanish nouns (http://memory.psych.upenn.edu/WordPools (*33*)). Word presentation is 1600 ms with a jittered 750-1200 ms (randomly sampled uniform distribution) blank screen after each word. After encoding is the distractor period, where participants performed 20 seconds of arithmetic math problems to disrupt their memory for recently-shown items. The problems were of the form A+B+C=??, where each letter corresponds to a random integer and participants typed their responses on the laptop keyboard. The final phase is retrieval, which began with a series of asterisks accompanied by a 300 ms, 60 Hz tone to signal for the participants to begin recalling as many words from the most recent list–in any order–they could remember for 30 seconds. Their vocalizations were recorded and later annotated offline using Penn TotalRecall (http://memory.psych.upenn.edu/TotalRecall) to determine correct recalls. For each session the participant began with a practice list of the same words that is not analyzed. The FR dataset includes 180 participants.

We group trials in the free recall task based on a number of criteria. 1st recalls refer to the first correct recall of a list in a given retrieval period. ≥2nd recalls refer to all correct recalls after the first correct recall if any occurred for that list. Intrusions include words said by the participant that were either from a previous list or were not from any list.

### Categorized free recall task (catFR)

A second variant of delayed free recall that participants performed is identical to FR except that the words in each list were presented in pairs that were semantically related (**Fig. 4a**). A word pool with 25 categories of 12 semantically-related words was created using Amazon Mechanical Turk to crowd source typical exemplars for each category (*33*) and was used for every session. For the creation of each 12-word list, 3 categories were randomly selected, and words were presented in sequential pairs but with no two pairs from the same category presented back-to-back. This setup allowed us to study both adjacently (same pair) and remotely presented words from the same category. Once again, each session began with a practice list of the same (unrelated) words that is not analyzed. The catFR dataset includes 104 participants with 65 of those participants also those that contributed to FR.

### Intracranial electroencephalogram (iEEG) recordings

iEEG was recorded from subdural grids and strips (intercontact spacing 10.0 mm) or depth electrodes (intercontact spacing 2.2-10.0 mm) using DeltaMed XlTek (Natus), Grass Telefactor, Nihon-Kohden, or custom Medtronic EEG systems. Signals were sampled at 500, 512, 1000, 1024, 1600, 2000 or 2048 Hz but downsampled using a Fourier transformation to 500 Hz for all analyses except for the control ripple detection using >250 Hz IED detection, where we downsampled to 1000 Hz and removed participants recorded at *<*1000 Hz. Initial recordings were referenced to a common contact, the scalp. or the mastoid process, but to eliminate potentially system-wide artifacts or noise and to better sense ripples locally we applied bipolar rereferencing between pairs of neighboring contacts. Bipolar referencing is ideal as the spatial scale of ripples is unlikely to exceed the inter-electrode spacing (*10*), thereby allowing us to localize ripples by eliminating system or muscle artifacts common to neighboring electrodes. Line removal is performed between 58-62 and 178-182 Hz using a 4th order Butterworth filter (120 Hz is in our sensitive ripple range and we did not find artifacts in these frequencies).

### Ripple detection

We utilized an algorithm recently shown to isolate ripples in human hippocampus (*7*) that is based on sharp wave ripple detection in rodents (*30*) and interictal epileptiform discharge (IED) removal in epileptic participants (*37*). The local field potential (LFP) from bipolar iEEG channels is filtered using a 70-178 Hz bandpass linear-phase Hamming-windowed FIR filter with a transition width of 5 Hz. The analytic signal envelope is then calculated using a Hilbert transform and extreme values were clipped to 4 SD to eliminate biasing due to extreme amplitudes (*30*). The resultant signal is then squared, smoothed with a 40 Hz low-pass Kaiser-windowed FIR filter with a transition width of 5 Hz, and the mean and SD were computed across all recalls in a session for that bipolar channel to set a ripple detection threshold. We then defined candidate events as points where the unclipped, squared envelope exceeded 4 SD above the mean. Each candidate event is expanded until the envelope fell below 2 SD and is considered a ripple if it is longer than 20 ms, shorter than 200 ms, and didn’t occur within 30 ms of another ripple, in which case they were merged. Finally, ripples within 50 ms of an IED were removed to avoid potentially pathological events (*7, 8*). IED detection followed a similar algorithm as outlined for ripples (*7, 37*): LFP is filtered using a 25-58 Hz bandpass linear-phase Hamming-windowed FIR filter with a transition width of 5 Hz, Hilbert-transformed, squared, 40 Hz low-pass filtered, and events 4 SD above the mean across recalls for that channel were considered IEDs.

Ripples were treated as discrete events throughout the paper (**Fig. 1c, 2a, etc**.) set to the beginning of each detected ripple, although since most ripples were between 20-40 ms in duration (**Supplementary Fig. 1a**) differences in timing relative to behavior were small. The ripple detection algorithm yields events that peak ~90 Hz, similar to recent work (*7–10*). Notably, using a different algorithm shown to detect ripples in hippocampus (*8*) and MTL (*10*) yields similar spectrograms and confirmed our main findings (**Supplementary Fig. 5**). Similarly, utilizing the primary algorithm with a higher detection range (125-200 Hz), in an attempt to isolate ripple events from sharp wave ripple–associated fast gamma (*29*) once again confirmed our main findings with events that peaked ~150 Hz (**Supplementary Fig. 6**).

Most participants had multiple MTL contacts within their montage, thereby providing multiple channels of iEEG for every recall. As has been done in past work (*7, 10*), since the spacing of clinical electrodes (2-10 mm) is much farther than ripples are expected to travel in the brain (*<*0.2mm, (*29*)), we treated each recalled word for each channel as a separate “trial”. Still, to ensure there is no redundancy across channels, we took three steps to guard against ripples being counted more than once.

First, to investigate ripples aligned to the time of recall for each participant, we had to account for any recalls that came close in time to a previous recall. Therefore, if a participant recalled a word within 2 s of a previous recall the latter recall is removed from our sample. This process allowed us to create peri-vocalization time histograms (PVTHs) aligned ±2 s to recall throughout the paper without double-counting ripples.

Second, after detecting ripples on each channel for a given participant, we created PVTHs (10 ms bins) for each channel for each region (i.e. hippocampus, entorhinal cortex, and parahippocampal cortex). Then we measured the pairwise correlations between the average PVTHs for each channel. For example, if a participant had 4 hippocampal channels, we would measure 6 correlations. We then averaged these correlations and if they were ≥0.2 we removed this session. We decided upon a 0.2 threshold based on 5 test participants during initial development of our algorithms, and as shown in **Supplementary Fig. 2a**, for the FR dataset most sessions fell within a normal distribution below 0.2.

Third, we manually inspected the raster plots (**e.g. Fig. 2a & Fig. 4b**) for every participant to ensure there is no redundancy across electrodes. Since ripples are fairly low-frequency events (~0.25-0.50 Hz, see PVTHs) it is clear if two channels had consistently overlapping ripples since only 1-2 ripples would occur in the 4 seconds surrounding each recall. Since as explained above ripples should not volume conduce across the intercontact spacing the occasional pair of channels with overlapping ripples were exclusively those where two bipolar pairs shared the same contact. In other words, if the first bipolar pair was LA1-LA2 and the second pair was LA2-LA3 the same ripples might show up in each channel. This is likely due to the ripples being localized very close to the shared (LA2, in this example) contact, thereby showing in both electrode pairs after subtracting background voltages in the surrounding pairs (LA1 and LA3). In the end, as is clear from the final raster plots **Figs. 2a & 4b**, there is no indication of redundant signals within the same participant after these series of checks.

There were also three steps we took to eliminate bad sessions from our sample. First, we only kept bipolar pairs with average ripples rates across recalls for the session between 0.1-1.0 Hz. The resulting distribution of ripples rates for FR is shown in **Supplementary Fig. 2b**. Second, for each channel in each session we measured the trial-by-trial correlation (using 10 ms bins, as above) and removed channels >0.05 to ensure there were not consistent artifacts across recalls. Indeed, the correlations across trials within a session rarely deviated beyond ±0.025 **Supplementary Fig. 2c**. Third, we manually eliminated channels with bad electrodes as identified by the clinicians at the hospitals or those with significant noise artifacts as detected by manual inspection of PVTHs. *<*5% of channels were removed via the combination of these steps.

Finally, we ran a control analysis where all electrodes in clinically-identified seizure onset zones (SOZs) were compared to those electrodes not identified as in SOZs. participants without reports (*<*20%) were omitted from this analysis.

### Anatomical localization

Pre-implant structural T1- and T2-weighted MRI scans were used to define the anatomical regions for each participant in addition to a post-implant CT scan to localize electrodes in the participant brain. Electrode contacts were semi-automatically localized using Voxtool (https://github.com/pennmem/voxTool) and the MRI and CT scans were coregistered using Advanced Normalization Tools (*23*) to align the brain regions to the electrode montage. Bipolar electrode pairs in hippocampal subfields CA1 and DG were localized using a combination of neuroradiologist labels (Joel M. Stein, Penn Medicine) and an automated segmentation process utilizing the T2 scan (*22*). To localize electrode pairs in entorhinal and parahippocampal cortex, we used a combination of neuroradiologist labels and an automated segmentation pipeline combining whole-brain cortical reconstructions from the T1 scan in Freesurfer (*38*), an energy minimization algorithm to snap electrodes to the cortical surface (*39*), along with boundaries and labels from the Desikan-Killiany-Tourville cortical parcellation protocol (*24, 40*). The point source of iEEG for bipolar electrode pairs was considered to be the midpoint between adjacent electrode contacts, except for when using a stricter definition where both electrode contacts were also required to be in the region as well (Supplementary Fig. 3). Exposed electrode contacts were typically 1-2 mm in diameter and 1.4-2.5 mm in length with the smallest contacts 0.8 mm in diameter and 1.4 mm in length.

### Semantic and temporal clustering

To study contextual reinstatement we investigated the clustering between contiguous recall transitions, in which we expect participants to recall words with semantic or temporal relationships to the previous word they recalled. For semantic clustering, we focused on the catFR dataset, as this task was specifically designed for participants to remember words drawn from semantic categories. As explained in the catFR section above, each 12-word list in this task had words drawn from 3 categories, with the 4 words from the same category presented in non-contiguous pairs. This setup provided a 2×2 matrix of possible transitions between consecutive, correct recalls: semantic vs. non-semantically related words (i.e. from the same category or not) and adjacent vs. remote words (i.e. words shown back-to-back during the encoding phase or not). This setup and the resulting proportions of transitions for adjacent semantic, remote semantic, and remote unclustered (non-semantic) are shown in **Fig. 4a**. Adjacent, unclustered transitions were only 3% of recalls so were excluded from analysis. The remaining recalls were those that led to intrusions (12%) or the last word recalled from that list (28%) which therefore had no transition. For each comparison between high- and low-clustering in **Fig. 4h-i** the groups of trials that contributed to the PVTHs were pooled together across participants using the given categories and the ripple rates were measured for each from −600 to −100 ms before recall. Many of the PVTHs show a rise in ripples well before −600 ms (**Fig. 4h-i & Supplementary Fig. 8a**), and the effect sizes were typically larger when using a time range −1100 to −100 ms before recall, but for consistency with the rest of the paper and our pre-registration we used only the −600 to −100 ms time range.

We also tested temporal transitions using the FR dataset, which utilized 12 unrelated words per list making it more conducive to isolating temporal clustering. Here for each transition between correct recalls we found the distance (lag) from those two words on the list. If the words were presented back-to-back on the list (either forwards or backwards) the absolute lag is 1. We compared lag=1 recalls to absolute lag≥4 recalls as done in previous work (*35*) to compare high- and low-temporal clustering.

Recalls that began with the first item shown during the encoding period and led to a chain of lag=1 transitions were removed from analysis for both FR and catFR until the recalled word was lag¿1. For example, if a person recalled the 1st, 2nd, 3rd, and 5th words that were presented during encoding, and in that same order, the first three of those recalls would be removed as they are likely due to a rehearsal strategy and not free recall (the last would be kept). However, due to the presence of the arithmetic distractor, recalls of this form were rare.

### Plots

Raster plots were formed by taking the time of each recall and plotting the time of the beginning of each detected ripple. As explained in Ripple Detection, any recalls within 2 s of a previous recall were removed from consideration in order to avoid double-counting ripples. Therefore, every ripple in the raster and PVTHs are unique events. PVTHs were formed by binning ripples (100 ms bins) and averaging the raster plots across participants after separating recalls into two groups: the first correct recall from each list or the remaining correct recalls from each list (≥2nd). For visualization only, these PVTHs are shown triangle smoothed using a 5-bin window (*7*) and a separate linear mixed model with sessions nested in participants is run at each bin to calculate the error bars (SE). Ripple rates are the frequency in Hz. within each bin. Dashed light gray horizontal lines (set to 0.35 Hz for the main figures) for all PVTHs are there to serve as visual aids for comparison between figures.

In order to visualize the consistency of the **PRE** across participants we fit a separate linear mixed effects model for each participant (Eq. 2) and plotted the t-scores for each given region (**Figs. 2c, 3e-f, & 4e**). We only plotted participants with ≥20 trials of ≥2nd recalls and minimum ripple rates of at least 0.1 Hz for both of the input bins in 2.

To compare **PRE** between regions **Fig. 3g** we show pairwise comparisons of t-scores for those participants with electrodes in at least two of our regions under study. For each bar only participants with ≥25 trials for both regions being compared were included in these plots. The sample sizes of participants for each bar between the pre-recall and post-recall tests (**Figs. 3g-h**) are slightly different based on whether the mixed effects model (4) converged, as some participants had relatively sparse ripples rates.

### Held out data and pre-registration

The unparalleled size of our datasets (the FR dataset alone has 20x more trials than previous studies of ripples in humans (*7, 8, 10*)) permitted us to come up with our initial hypotheses based on analysis of only ~40% of the FR and catFR datasets. That is, after creating the raster plots and ensuring all data was in usable form via the data-cleaning steps outlined above, we used a random kernel to select a subset of participants that had 40% of hippocampal trials for FR. We performed the same steps for catFR using a separate randomization to select a subset of participants that had 40% of hippocampal trials. Since the 40/60% partition was based on trials in the entirety of the hippocampus, the held out participants did not necessarily have 60% of data for the individual subfields CA1 or DG, nor did they necessarily have 60% of data for entorhinal or parahippocampal cortex. However, we expected by chance to have at least close to an ~50% partition between the exploratory and confirmatory sets for all tests. The number of trials or participants that go into each analysis are labeled on each plot or within the caption. We registered these hypotheses on the Open Science Framework (https://osf.io/y5zwt), which contains specific details on our randomization and shows initial figures for most analyses presented in this paper done with only the exploratory ~50% of the data. Throughout the paper, significance is assessed on the whole dataset.

### Equations

Linear mixed effects models were run using the function MixedLM in the python package statsmodels with restricted maximum likelihood and Nelder-Mead optimization with a maximum of 2000 iterations. The following equations are written in pseudocode of the inputs to statsmodels. Statistics are presented as: *β* ± *SE, p* − *value*, where *β* is the coefficient being tested in Equations 1-5 and *SE* is the standard error.

To test the hypothesis that the **PRE** is stronger for recalls that were not the first from each list (**Fig. 2b**), we used the linear mixed effects model:

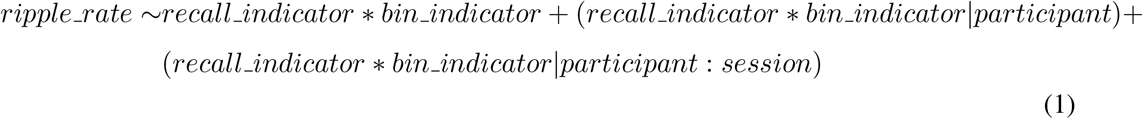

where *recall_indicator* is 0 for 1st recalls from each list and 1 for ≥2nd recalls from each list, *bin_indicator* is 0 for the bin −1600 to −1100 ms and 1 for the bin −600 to −100 ms aligned to time of recall, (*recall_indicator* ∗ *bin_indicator*|*participant*) are random intercepts and slopes for each factor and the interaction in each participant, (*recall_indicator* ∗ *bin_indicator*|*participant* : *session*) are random intercepts and slopes for sessions nested in each participant, and *ripple_rate* is the ripple rate for each trial. The null hypothesis is no difference in the interaction between recall type (1st vs. ≥2nd) and bin.

We also investigated **PRE** individually for each participant (**Fig. 2c**). We fit a linear mixed model on the participant’s ≥2nd recall trials:

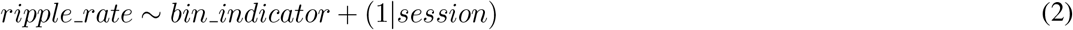

where 1|*session* is a random intercept for different sessions and the other factors are the same as in Eq. (1). The null hypothesis is no difference in ripple rate between the −600 to −100 ms bin and the same bin aligned 1 second earlier.

Similarly, we assessed **PRE** for individual electrode pairs within participants so we could overlay t-scores on their anatomical localizations (**Fig. 1e & 4f**). These t-scores came from the coefficient fit for *bin_indicator* in Eq. (2) with trials selected only for single electrode pairs at a time.

To test the hypothesis that participants with better memories show a stronger **PRE** (reported in the caption of **Fig. 2d**), we used the linear mixed effects model:

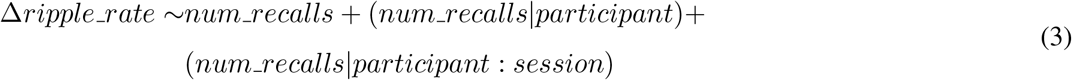

where *num_recalls* is the number of total recalls by the participant from the list the trial came from, Δ*ripple_rate* is the difference in ripples from the bin −600 to −100 ms and −1600 to −1100 ms, and the other factors are random effects for participant and session nested in participant as in Eq. (1). The null hypothesis is no difference between number of recalls per list and change in ripple rate.

To make pairwise comparisons between regions to test if some regions have a stronger **PRE** than others (**Fig. 3g**), we used the linear mixed effects model:

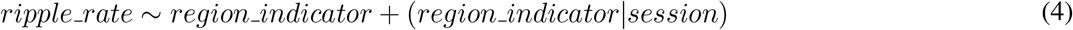

where *region_indicator* is 0 or 1 for two different regions (in the order shown beneath each swarm plot in **Fig. 3g**), (*region_indicator*|*session*) is a random intercept and slope for each session, and *ripple_rate* is the ripple rate in the bin from −600 to −100 ms aligned to recall. Note that every test is for bipolar electrode pairs in different regions for the same participant, therefore only variance across sessions had to be accounted for. The null hypothesis is no difference in **PRE** between regions. Significance for each of the 6 pairwise comparisons is assessed with an FDR-corrected (Benjamini-Hochberg) t-test to correct for the 6 comparisons.

We also made pairwise comparisons between regions for post-recall ripples (**Fig. 3h**). The equation is the same as Eq. (4), except the ripple rates were from the bin 200 to 700 ms after recall.

Finally, to compare **PRE** between two groups of trials, i.e. high- vs. low-clustering trials (**Fig. 4h-j**), correct vs. intrusion trials (**Fig. 2e**), or SOZ vs. non-SOZ (**Supplementary Fig. 7**), we used the linear mixed effects model:

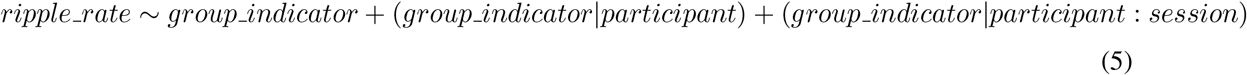

where *group_indicator* is 0 for the first group of trials and 1 for the second group of trials, (*group_indicator*|*participant*) is a random intercept and slope for each participant, (*group_indicator*|*participant* : *session*) is a random intercept and slope for each session nested in participants, and *ripple_rate* is the ripple rate in the bin from −600 to −100 ms aligned to recall. The null hypothesis is no difference in **PRE** between the two groups. Negative coefficients indicate a decrease in ripples between the first and second group (e.g., a drop from high- to low-clustering).

## Supplementary figures

**Supplementary Fig. 1.**
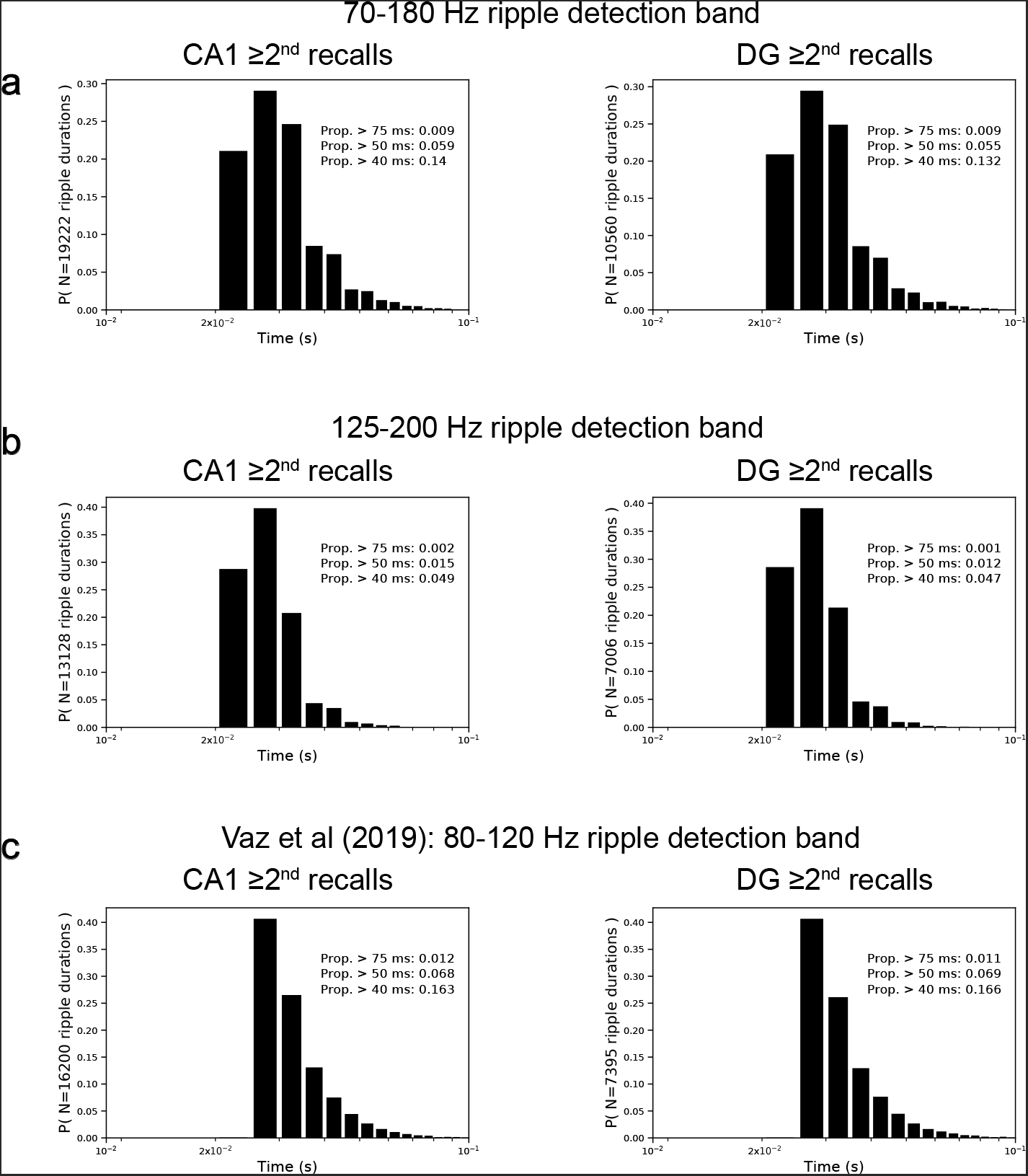
Durations of ripples. **a**, The distribution of ripple durations for the detection algorithm used throughout the main text for all ripples contributing to the ≥2nd recall PVTHs in Fig. 2b. **b**, The distribution of ripple durations for the same ripple detection algorithm using only the higher frequency range (125-200 Hz). **c**, The distribution of ripple durations for the Vaz et al. ripple detection algorithm (*10*).

**Supplementary Fig. 2.**
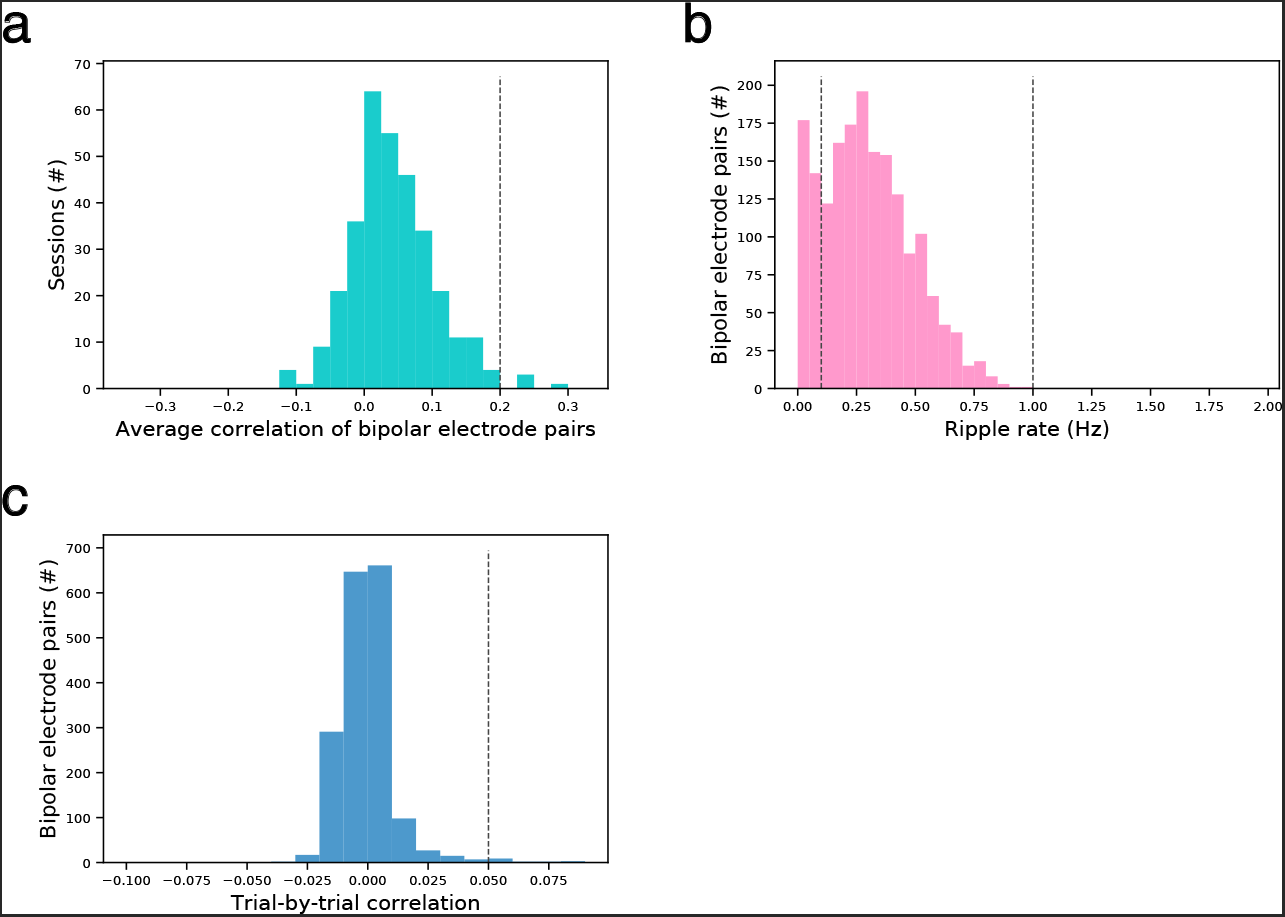
Distribution of parameters used to remove bipolar pairs or sessions. **a**, Average pairwise correlations between ripple rate PVTHs within each session for participants with multiple hippocampal bipolar pairs. Sessions were removed if the pairwise correlation was ≥0.2 (dotted line), which would indicate artifacts across channels. **b**, Average ripple rate at each hippocampal bipolar pair for each session. Bipolar pairs with ripple rates *<*0.1 or >1.0 Hz. (dotted lines) were removed, which would indicate pairs likely not in hippocampus or with noise artifacts, respectively. **c**, Average pairwise correlations between ripple rate PVTHs within each hippocampal bipolar pair for single sessions. Bipolar pairs were removed for a given session if the correlation was ≥0.05 (dotted line), which would indicate artifacts across trials.

**Supplementary Fig. 3.**
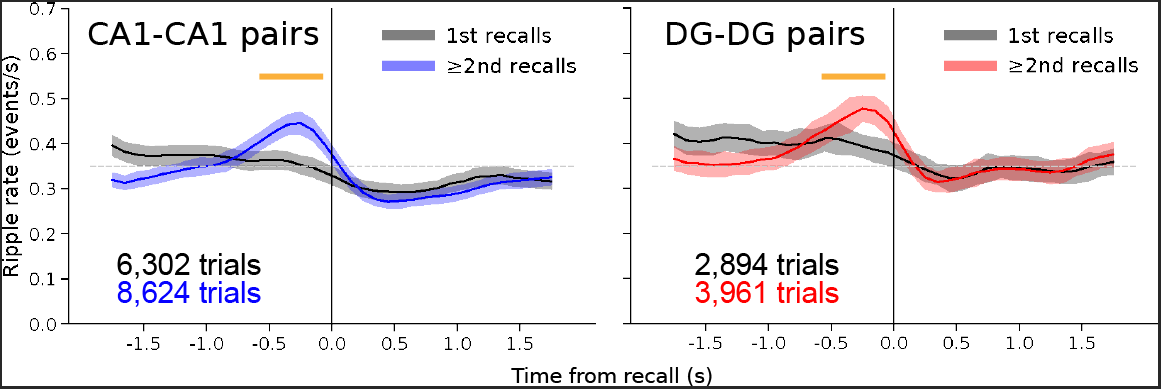
Stricter localization of bipolar pairs in CA1 and DG still shows a significant PRE in each subfield. PVTHs of ripples aligned to recall for only those bipolar pairs where both contacts are individually localized to CA1 (CA1-CA1, left) or both contacts are individually localized to DG (DG-DG, right). CA1: 175 sessions from 82 participants. DG: 101 sessions from 46 participants. The **PRE** is significant for both CA1-CA1 pairs (*β* = 0.11±0.030, *P* = 1.9 × 10^−4^, Eq. 1) and DG-DG pairs (*β* = 0.12±0.032, *P* = 2.4 × 10^−4^, Eq. 1).

**Supplementary Fig. 4.**
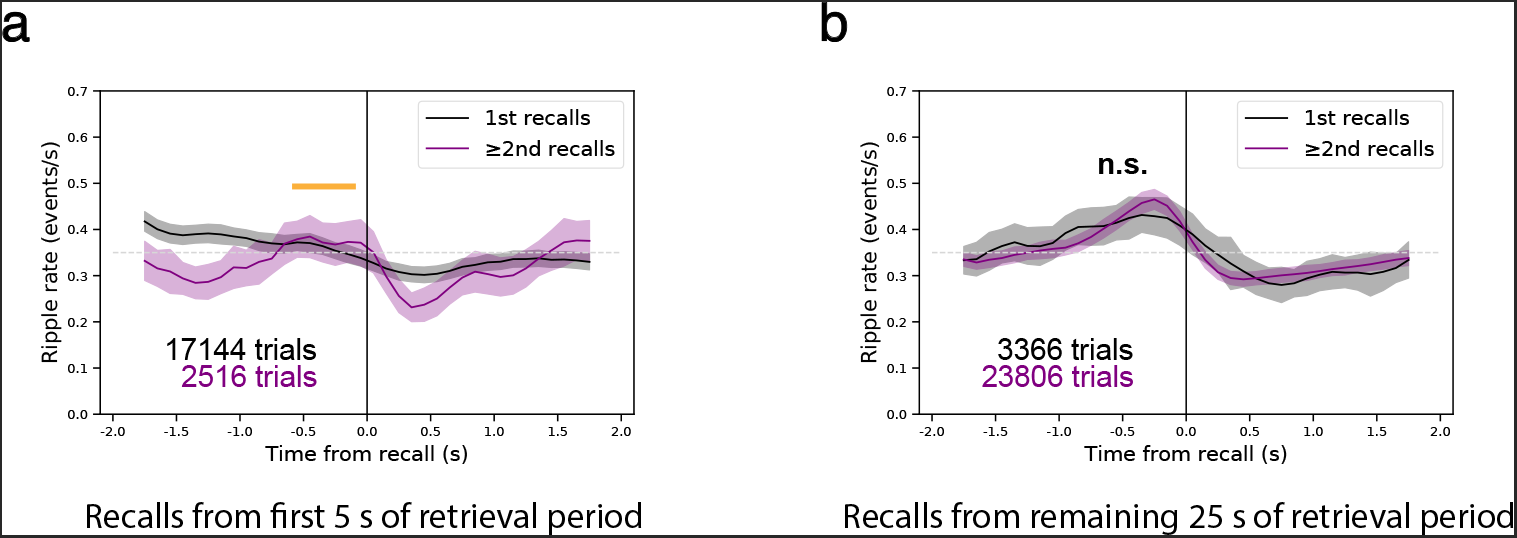
PVTHs of ripples aligned to recall for less than or greater than 5 s of retrieval period. **a**, Ripples aligned to time of recall for recalls that occurred in the first 5 s of the retrieval period. Trials are combined for CA1 and DG to increase the sample size of ≥2nd recalls and due to their nearly identical effects in **Fig. 2b**. The **PRE** is significant ≥2nd trials (*P* = 0.0018, Eq. 1). **b**, Same, but for recalls occurring in the final 25 s of the retrieval period. There is no significant difference in **PRE** (*P* = 0.35, Eq. 1).

**Supplementary Fig. 5.**
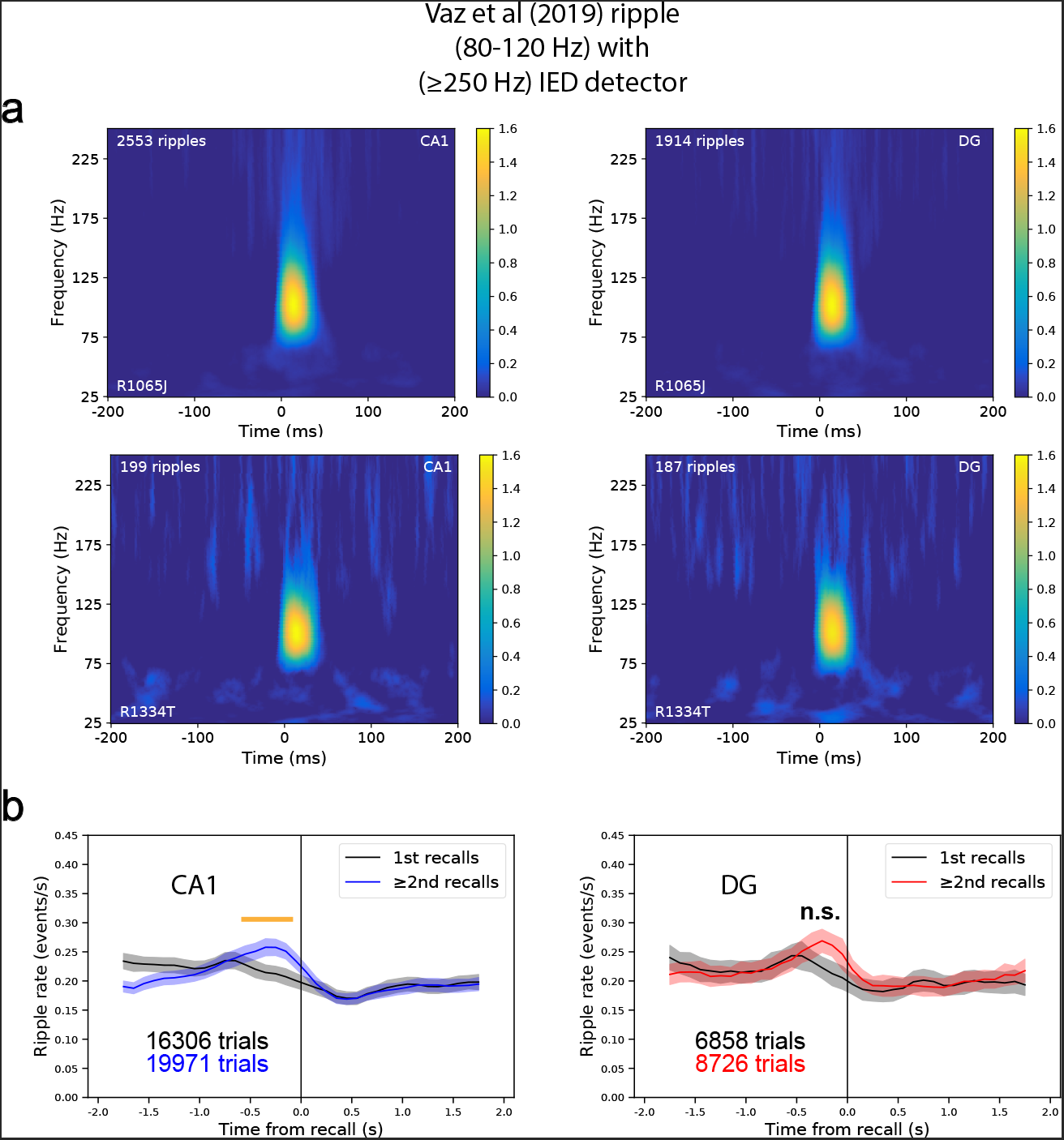
PRE still present using detection method from Vaz et al (2019). **a**, Average spectrograms across ripples for same two participants shown in Fig. 1D for both CA1 and DG. Note the peak is still ~90 Hz as in the original method. **b**, PVTHs for 1st vs. ≥2nd recalls for CA1 and DG using the Vaz et al. detector. The **PRE** was significantly greater for ≥2nd recalls for CA1 (*P* = 2.5 × 10^−4^, Eq. 1) but not for DG (*P* = 0.13).

**Supplementary Fig. 6.**
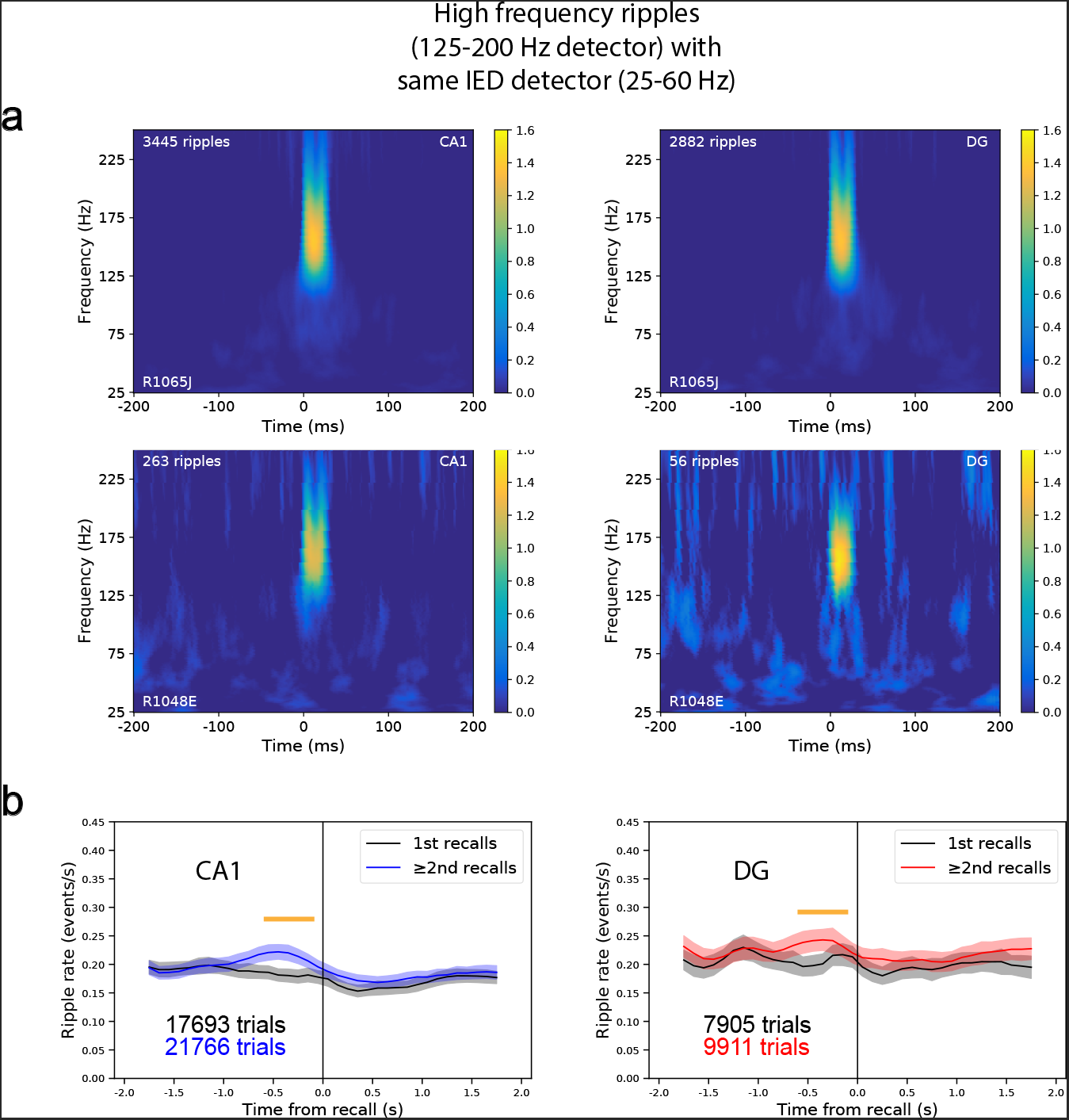
PRE still present using higher frequency (125-200 Hz.) ripple detection method. **a**, Average spectrograms across ripples for same two participants shown in Fig. 1D for both CA1 and DG. Note the peak ~150 Hz. **b**, PVTHs for 1st vs. ≥2nd recalls for CA1 and DG using high frequency detector. The **PRE** is significantly greater for ≥2nd recalls for CA1 (*P* = 2.2 × 10^−4^, Eq. 1) and DG (*P* = 0.019).

**Supplementary Fig. 7.**
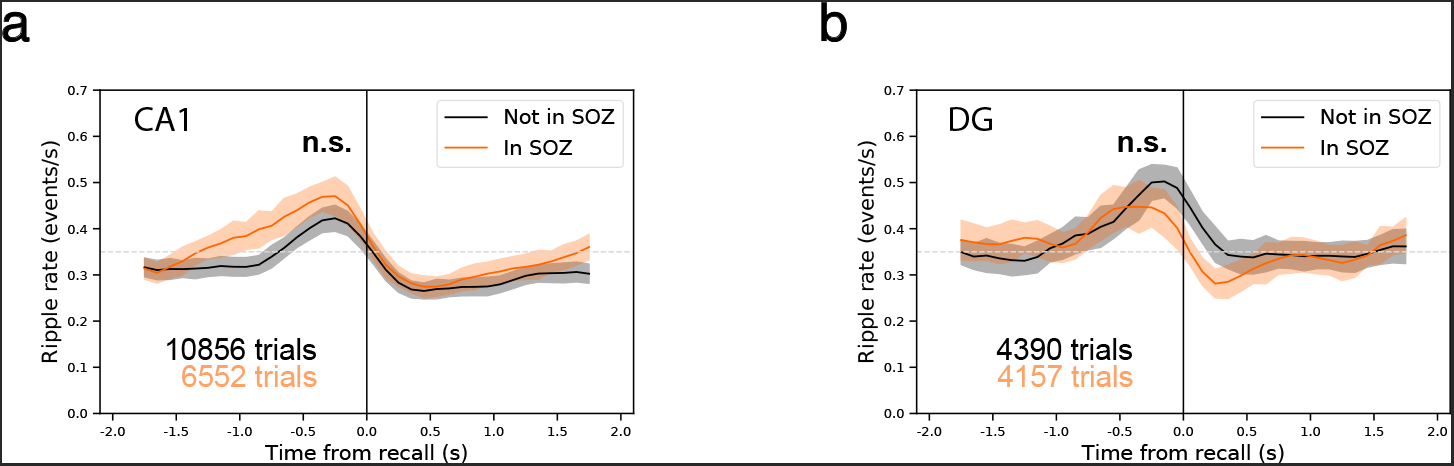
PRE occurs both within and outside seizure onset zone (SOZ). **a**, For all participants that had a clinically-defined seizure onset zone, we plotted PVTHs aligned to recall for all CA1 trials recorded in and not in the SOZ (participants with no SOZ information were excluded) for ≥2nd recalls. Both trials in SOZ (*P* = 1.6 × 10^−13^, Eq. 2) and not in SOZ (*P* = 1.6 × 10^−16^, Eq. 2) showed a significant **PRE**. However, **PRE** between these groups of trials is not significantly different (*P* = 0.40, Eq. 5). **b**, Same as **a**, but for electrodes in DG. Both trials in SOZ (*P* = 9.2 × 10^−4^, Eq. 2) and not in SOZ (*P* = 7.0 × 10^−11^, Eq. 2) showed a significant **PRE. PRE** between these groups of trials is not significantly different (*P* = 0.071, Eq. 5).

**Supplementary Fig. 8.**
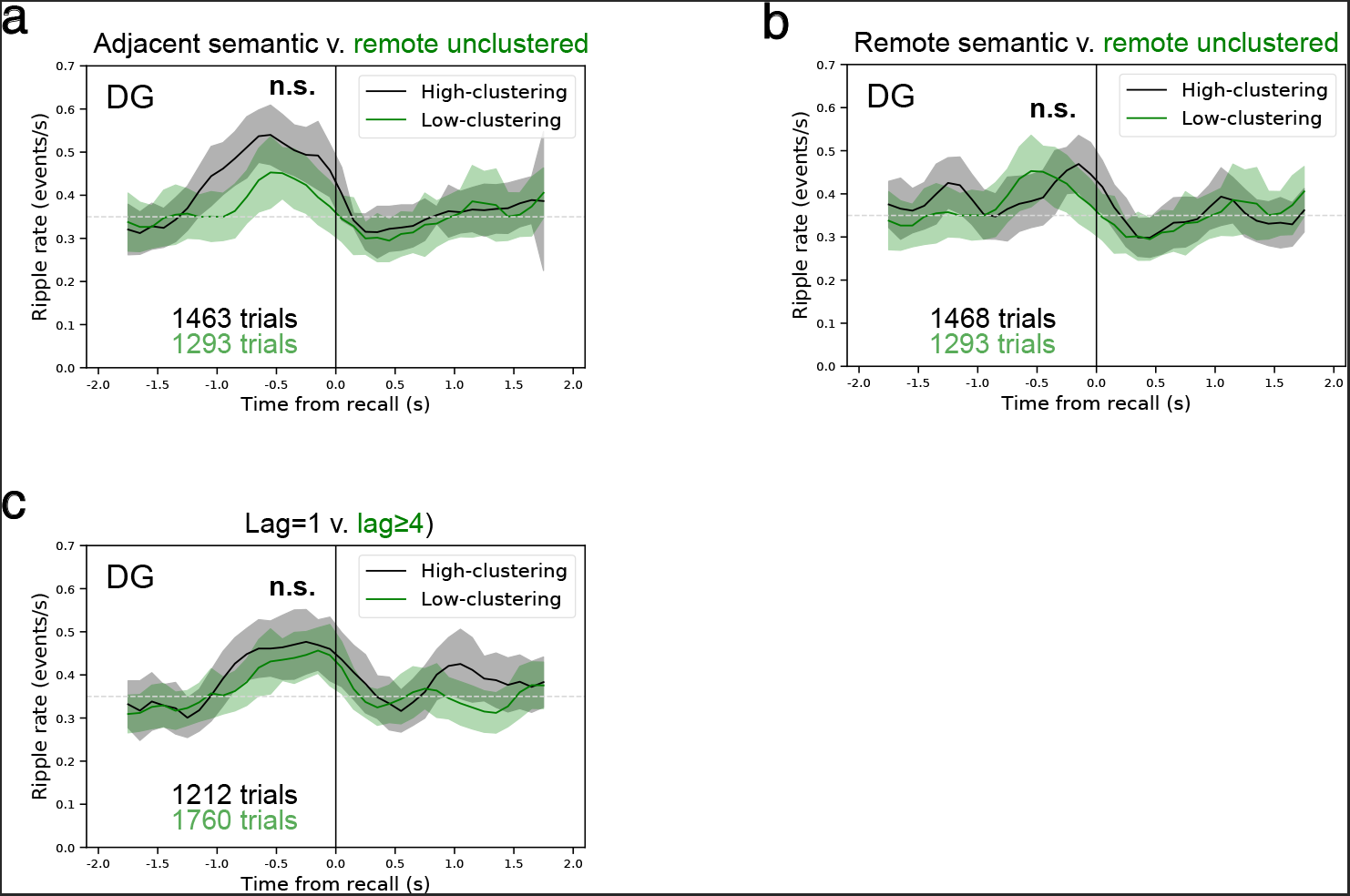
Contextual reinstatement and the PRE in dentate gyrus. **a**, PVTH of catFR trials comparing adjacent semantic vs. remote unclustered trials, a test of contextual reinstatement vs. no contextual reinstatement. Significance of coefficient comparing each trial type in mixed model (Eq. 5): held out data: DG, *P* = 0.054; 100% of data: DG, *P* = 0.022 (FDR-corrected across 6 tests of Eq. 5, **Figs. 4h-j & Supplementary Fig. 8a-c**). **b**, PVTH of catFR trials comparing remote semantic vs. remote unclustered trials, a test of semantic reinstatement vs. no contextual reinstatement. Same significance test and conventions as **a**: held out data: DG, *P* = 0.95; 100% of data: DG, *P* = 0.56 (FDR-corrected). **c**, PVTH of FR data (original task outlined in **Figs. 1-3** comparing adjacent recalls (lag = 1) vs. remote recalls (lag≥4), a test of temporal reinstatement. Same significance test and conventions as **a**: held out data: DG, *P* = 0.87; 100% of data: DG, *P* = 0.39 (FDR-corrected).

